# Comparative analysis of cloacal microbiota in *Henophidia* (non-venomous) and *Caenophidia* (venomous) snakes

**DOI:** 10.64898/2026.05.13.724777

**Authors:** Ehsan Ghasemian, Sedreh Nassirnia, Trestan Pillonel, Simon Rüegg, Sebastien Aeby, Claire Bertelli, Nicole Borel, Gilbert Greub

## Abstract

The evolutionary divergence between *Henophidia* (non-venomous) and *Caenophidia* (venomous) snakes has produced distinct cranial morphologies, digestive strategies, and presence of specialised venom systems in *Caenophidia*, yet the extent to which these long-standing diverging trajectories have shaped cloacal microbiota assembly remains poorly understood. We characterised cloacal microbiota in 70 captive snakes (52 *Caenophidia*, 18 *Henophidia*) by 16S rRNA amplicon sequencing. Beta diversity was tested by PERMANOVA, differential abundance by ANCOM-BC2, community types by Dirichlet Multinomial Mixture modelling (DMM), and microbial interactions by SparCC co-occurrence networks. Predicted functional potential (PICRUSt2) was analysed by ALDEx2 differential abundance testing and elastic net feature selection. *Henophidia* exhibited significantly higher bacterial richness and greater compositional variability than *Caenophidia*. Community composition showed clade-associated differences (PERMANOVA) and partitioned into two distinct DMM community types. The *Henophidia* network was 11.9-fold denser and more modular, with *Burkholderiaceae* as a keystone hub, whereas the *Caenophidia* network was sparse. *Henophidia* showed predicted enrichment in C1 metabolic pathways (ethylmalonyl-CoA, formaldehyde assimilation I, glycine betaine degradation I, methylaspartate cycle), aromatic compound catabolism, and nitrogen recycling, whilst *Caenophidia* showed enrichment in allantoin and glucuronate degradation. This multi-method analysis suggests *Burkholderiaceae* as a candidate keystone taxon in *Henophidia* and indicates that phylogenetic clade is a major contributor to cloacal microbiota structure. The lower richness in *Caenophidia* raises a testable hypothesis that broad-spectrum antimicrobial activity of their venom components may selectively filter susceptible microbial lineages, motivating future shotgun metagenomic studies in wild populations of snakes.

## Introduction

Snakes represent one of the most successful vertebrate evolutionary lineages, comprising more than 4,000 species distributed across a wide range of ecological niches worldwide [1]. Within this diversity, the phylogenetic split between Henophidia and *Caenophidia* constitutes a fundamental evolutionary divergence that occurred approximately 60-70 million years ago, giving rise to major morphological, physiological, and ecological differences [2]. Henophidia includes families such as *Boidae*, *Pythonidae*, and *Xenopeltidae*, which are characterised by vestigial pelvic girdles, broader cranial structures, and predominantly constriction-based prey capture strategies. In contrast, *Caenophidia* encompasses the majority of existing snake families, including *Colubridae*, *Viperidae*, and *Elapidae*, and exhibits more derived cranial kinesis, specialised dentition, and a wider range of prey control mechanisms [2]. Henophidian species typically consume larger prey relative to body size and display prolonged digestive periods, whereas caenophidian species show greater dietary diversity and more rapid digestive processing [3].

The cloacal microbiota of snakes exhibits several physiological functions, including nutrient metabolism, modulation of the immune system, exclusion of pathogens, and maintenance of gut homeostasis [4,5]. Unlike mammals, reptilian digestive systems exhibit distinctive characteristics such as irregular feeding patterns, a high tolerance to prolonged fasting, and pronounced post-prandial organ remodelling, all of which strongly influence host-microbiome interactions [6,7]. The cloaca, which serves as the terminal chamber for the digestive, urinary, and reproductive tracts, constitutes an ecological niche in which microbial communities interact simultaneously with multiple physiological systems. Previous studies have shown that gut microbial composition in snakes is shaped by host phylogeny, diet, habitat, and environmental factors, with dominant phyla typically including *Bacteroidetes*, *Firmicutes*, *Proteobacteria*, and *Fusobacteria* [8,9].

A major characteristic distinguishing most caenophidian from henophidian species is the presence of venom systems, which play roles in prey immobilisation and likely digestion [10–13]. Venom composition varies considerably amongst caenophidian families, with vipers possessing predominantly cytotoxic and haemorrhagic toxins, elapids producing neurotoxic compounds, and colubrids exhibiting diverse venom profiles [14]. Importantly, snake venoms contain numerous enzymatic components, including metalloproteases, serine proteases, phospholipases, and hyaluronidases, which can facilitate the pre-digestion of prey tissues and may, in turn, influence subsequent microbial colonisation and metabolic activity within the gastrointestinal tract [11,12,15–17]. Prior studies have shown that venom-mediated protein hydrolysis enhances digestive efficiency and nutrient availability, potentially creating distinct nutritional environments that favour specific microbial taxa [13,18]. Moreover, certain venom components possess antimicrobial properties that may selectively modulate gut and cloacal microbial communities, either through direct bactericidal effects or by altering local pH and biochemical conditions [15]. The absence of venom systems in henophidian snakes necessitates alternative digestive strategies, including prolonged gastric digestion and a greater reliance on endogenous digestive enzymes and microbial fermentation [19].

The differ ences in evolutionary trajectories, morphology, and functional ecology between *Henophidia* and *Caenophidia*, including variation in digestive physiology, feeding frequency, prey preferences, and venom systems, provide a valuable framework for investigating whether microbial communities in these two clades have evolved differently [20]. Phylogenetically conserved traits within each clade may impose distinct selective pressures on their microbiomes, potentially giving rise to clade-specific microbial signatures [21,22]. However, the extent to which long-standing evolutionary divergence has shaped cloacal microbiota remains poorly understood. Comparative analyses of cloacal microbial communities across these clades could therefore yield novel insights into the ecological success, dietary adaptations, and diversification patterns of these major snake lineages. To date, snake microbiome studies have predominantly focused on single species or limited taxonomic groups, constraining our ability to detect broad evolutionary patterns [7,23,24]. Here, using 16S rRNA amplicon sequencing and comparative analytical approaches, this study aims to characterise cloacal microbiota composition, diversity, and functional potential across a collection of henophidian and caenophidian snake samples obtained from private snake collectors in Switzerland.

## Materials and methods

### Sample collection

Five private collections (arbitrarily designated collections 1-5) contributed samples from 25, 7, 13, 15, and 10 exotic snakes, respectively. Cloacal and choanal swabs were collected from a selection of 137 snakes across these five collections, of which samples from 99 snakes were taken forward for DNA extraction according to Borel *et al*. [25] and microbiota profiling. This study was conducted within the framework of a *Chlamydiota* screening investigation in snakes, in which molecular diagnosis of *Chlamydiota* was the primary objective of DNA extraction [25–28]. Nevertheless, opportunistic sampling from this prospective cohort allowed us to explore the cloacal microbiota of caenophidian (venomous) and henophidian (non-venomous) snakes.

### 16S rRNA sequencing

Library preparation followed the Illumina 16S Metagenomic Sequencing Library Preparation protocol [29]. Briefly, the V3–V4 hypervariable region of the bacterial 16S rRNA gene was amplified in an initial 25-cycle PCR, yielding fragments of approximately 400 bp, after which sample-specific barcodes and Illumina adapters were incorporated in a subsequent 8-cycle indexing PCR. PCR products were assessed for fragment size and yield using a Fragment Analyzer with the NGS Fragment Kit (AATI, USA) and quantified using a Qubit fluorometer with the dsDNA High Sensitivity Kit (Thermo Fisher Scientific, USA). Libraries were normalised to 4 nM, pooled, and diluted to 10 pM, with 10% PhiX spike-in added as a sequencing control. Sequencing was performed on an Illumina NextSeq 1000 platform using the P1 v3 reagent kit, generating 2 × 150 bp paired-end reads. We included a defined mock community (MSA-2002™, ATCC, Manassas, USA) as a positive control to assess sequencing accuracy, an extraction blank as a negative control to detect reagent-level contamination introduced during DNA extraction, and a no-template PCR control to identify contamination arising during library preparation.

### Data processing

Raw 16S rRNA gene sequencing reads were processed using the in-house zAMP pipeline [30], which integrates quality assessment (FastQC), adapter trimming and read filtering (Cutadapt), error correction and amplicon sequence variant (ASV) inference (DADA2), and taxonomic classification against the Greengenes2 database.

Potential contaminant taxa were identified using the decontam package (v1.16.0) with the prevalence-based method (threshold = 0.1), comparing negative library controls (NLC) against true biological samples. To prevent over-removal of biologically relevant taxa, we applied a rescue strategy in which taxa flagged as contaminants were retained if they showed higher mean abundance in true samples than in negative controls. Taxa with true-to-NLC ratios ≥2 and higher prevalence in true samples were systematically rescued and kept in the dataset. Following decontamination, taxa and samples with zero abundance were pruned.

To account for uneven sequencing depth across samples, we employed a consensus rarefaction approach. All samples were rarefied to a uniform depth of 5,000 reads per sample. To minimise the stochastic effects inherent in single rarefaction events, we performed 100 independent rarefaction iterations, each using a unique random seed. Consensus abundances were then obtained by averaging taxon counts across all iterations, with final values rounded to integers. Twenty-nine samples were excluded due to insufficient sequencing depth, and the remaining 70 samples were rarefied and retained for downstream analyses. Sequencing generated 7,282,462 raw reads across the 70 samples included in the analysis, of which 5,571,674 were retained after quality filtering and 5,465,550 remained following chimera removal (Table S1).

### Alpha diversity analysis

Prior to fitting the alpha diversity models described below, the appropriate treatment of sex and husbandry facility as fixed or random effects was evaluated. Sex was retained as a fixed effect as factors with fewer than five levels yield unstable random-intercept variance estimates [31,32]. For husbandry facility (k = 5), a diagnostic protocol comprising intraclass correlation coefficients (ICC), boundary-corrected likelihood ratio tests using a 50:50 𝜒^2^ mixture [33], 𝛥AIC, singular-fit checks, and sensitivity analyses on the clade coefficient supported a fixed-effect specification: although ICCs were non-negligible for Chao1 (0.29) and Shannon (0.18), the corresponding likelihood ratio tests were non-significant (p = 0.17 and p = 0.21, respectively) with 𝛥AIC values falling within ±2, and Pielou’s evenness yielded a singular fit (ICC = 0). Combined with the small number of facilities (k = 5), which is at the lower threshold for stable random-effect estimation, and the strong confounding between clade and facility (Cramér’s V = 0.83, with two husbandries housing only a single clade), these results jointly supported modelling both sex and husbandry facility as fixed effects throughout the alpha diversity analyses.

To assess differences in alpha diversity metrics at ASV level, including richness (Chao1), diversity (Shannon index), and evenness (Pielou’s evenness) between *Caenophidia* and Henophidia clades, fitted linear models with clade as the main explanatory variable. Models were adjusted for potential confounders, including sex and husbandry facilities. Type II analysis of variance (ANOVA) was performed to evaluate the effect of clade while accounting for covariates. Statistical significance was determined at α = 0.05, and pairwise comparisons were conducted between clades for each diversity metric.

### Beta Diversity

Prior to the beta diversity analyses, the appropriate treatment of sex and husbandry facility as fixed or random effects was evaluated. Sex was retained as a fixed effect as factors with fewer than five levels are not amenable to random-effect treatment [31,32]. For husbandry facility (k = 5), a fixed-effect specification was supported by three considerations: first, the design comprised only five facilities, which is at the lower threshold for stable random-effect or strata-based estimation; second, clade and facility were strongly confounded (Cramér’s V = 0.83), with two of the five husbandries housing only a single clade, such that within-facility permutation strata would effectively reduce the test to comparisons among the three mixed - clade facilities (n = 47); third, PERMDISP indicated significant heterogeneity of dispersion across facilities for both Bray-Curtis (p = 0.002) and Jaccard (p = 0.006) distances, which is more transparently accommodated by including facility as an explicit covariate term than by restricting the permutation space.

ASVs were filtered using prevalence and detection thresholds (≥20% prevalence and ≥0.01% relative abundance) and subsequently aggregated to the genus level, an approach that was consistently applied across downstream analyses, including compositional profiling, differential abundance testing, community typing and co-occurrence network. Bray-Curtis and Jaccard distance matrices were then computed from the resulting genus-level abundance data using the phyloseq package. To test for significant compositional differences between clades whilst controlling for potential confounding factors, permutational multivariate analysis of variance (PERMANOVA) was performed using the adonis2 function from the vegan package with 999 permutations, including taxonomic clade, sex, and husbandry facility as explanatory variables. Type II sums of squares (marginal effects) were used to assess the effect of each variable while accounting for the others. To determine whether observed differences in community composition were driven by true compositional shifts (centroid location) or differences in within-group dispersion, multivariate homogeneity of group dispersions was tested using the betadisper function followed by permutation-based ANOVA (999 permutations). Non-metric Multidimensional Scaling (NMDS) ordinations was performed (k = 2, trymax = 100) to visualise community structure. Environmental fitting (envfit, 999 permutations) was used to project the 15 most abundant genera onto the NMDS ordination and assess their axis correlations.

### Constrained ordination

Canonical Analysis of Principal Coordinates (CAP) was performed using the capscale function from the vegan package to quantify the proportion of compositional variance explained by explanatory variables. Separate models were fitted for Bray-Curtis and Jaccard distance matrices, with taxonomic clade, sex, and husbandry facilities as constraining variables. Model significance was assessed using permutation tests (999 permutations). Type II (marginal) tests were performed using the *anova.cca* function to evaluate the independent contribution of each predictor and adjusted R^2^ values were calculated to account for model complexity relative to sample size. To identify taxonomic drivers of constrained variation, genus-level abundances were fitted as environmental vectors onto the ordination space using envfit function (999 permutations). Correlation strengths (r^2^) were used to assess the alignment of each genus with the constrained axes and the 15 genera with the highest r^2^ values were selected and overlaid to the ordination plots.

### Compositional profiling and core genera identification

Compositional profiles were summarised at the phylum and genus levels using stacked bar plots. Mean relative abundances were calculated per clade, with the top 10 phyla and top 25 genera displayed individually and all remaining taxa grouped as “Other”.

Core and clade-specific genera were identified based on within-clade prevalence and detection thresholds as described previously. Genera meeting these criteria in both clades were classified as core, whilst those detected exclusively in one clade were classified as clade-specific. Overlap and uniqueness between clades were then visualised using Venn diagrams.

### Differential Abundance Analysis

Differential abundance testing of bacterial genera between *Caenophidia* and *Henophidia* clades was performed using Analysis of Compositions of Microbiome with Bias Correction (ANCOM-BC2) in R. Reads aggregated to genus level were modelled with clade, sex, and husbandry facilities as fixed effects, using *Caenophidia* as reference. P-values were adjusted using Benjamini-Hochberg (BH) False Discovery Rate (FDR) correction (q < 0.05), and biologically meaningful differences were defined as an absolute log2FC > 1.

### Unsupervised community typing

Unsupervised community typing was performed using Dirichlet Multinomial Mixture (DMM) modelling on genus-level taxonomic profiles derived from the filtered dataset. DMM models were fitted for k = 1 to 8 community types, with the optimal k determined by minimising the Laplace criterion. Driver taxa for each community type were identified as the genera with fitted Dirichlet parameters (𝛼) above the 80^th^ percentile within that type; these parameters represent the genus-level weights of the corresponding multinomial mixture component and quantify each genus’s contribution to the expected composition of that community. To complement this, genera were ranked by log2FC in mean relative abundance between each community type and all other samples, with the top 30 per type reported as the most discriminatory taxa. Community composition and key taxa were visualised using stacked bar plots and dot plots, respectively. The associations between DMM community types and taxonomic clades were assessed using alluvial diagrams.

### Microbial Co-occurrence Network Analysis

Genus-level taxonomic profiles derived from the filtered dataset were used for co-occurrence network construction using the NetCoMi package. Networks were constructed separately for each clade using SparCC (Sparse Correlations for Compositional data) with 200 iterations to estimate correlations whilst accounting for the compositional nature of microbiome data and reducing spurious associations arising from the sum constraint. Network edges were defined by applying a correlation threshold of r ≥ 0.5 combined with local false discovery rate (lfdr) correction < 0.05 to identify statistically significant and strong associations. Network modules were identified independently within each clade-specific network using the fast greedy modularity optimisation algorithm, yielding network-specific module labels with no a priori cross-network correspondence. To quantify module-level overlap between clades, Jaccard similarity was calculated between every pair of modules from the two networks, excluding the cluster of unconnected nodes in each network. Hub genera were defined as nodes exceeding the 90^th^ percentile in at least one of the centrality metrics: degree, betweenness, closeness, and eigenvector centralities. Network topology was visualised by standard graph metrics, including number of nodes and edges, network density, clustering coefficient, modularity, and average path length. Statistical comparison of network structures between clades was performed using permutation-based testing (100 permutations).

### Functional analysis

To infer functional capacity from 16S rRNA gene amplicon data, ASV-level count tables were first filtered to retain taxa detected above 0.01% relative abundance in at least two samples within either clade, preserving clade-specific taxa whilst removing spurious low-abundance ASVs. Filtered ASV sequences and count tables were used as input to PICRUSt2 v2.5.2, which infers metagenome composition based on phylogenetic placement of marker gene sequences against a reference genome database. Predicted KEGG Ortholog (KO) abundances, Enzyme Commission (EC) number abundances, MetaCyc pathway abundances, and taxon-stratified KO contributions were generated with stratified output and per-sequence contribution tracking enabled. Samples with a weighted nearest sequenced taxon index (NSTI) > 0.2 were excluded from downstream analysis to ensure prediction reliability. Pathway abundances were further filtered to retain only those present in at least 60% of samples in at least one clade. Differential pathway abundance was assessed using two approaches. First, ALDEx2 was applied as a compositionally aware method employing Dirichlet Monte Carlo sampling, Welch’s t-test and BH correction. Second, bootstrap-stabilised elastic net regularised logistic regression (α = 0.5) was applied over 100 bootstrap iterations, retaining only features selected in ≥70% of bootstrap models as a stability criterion. Husbandry facilities and host sex were included as unpenalised covariates. For ALDEx2, significance was defined by combined criteria of adjusted p < 0.05, effect size ≥ 1.0, and centred log-ratio (CLR) difference > 2.0; for elastic net, pathways were retained if they showed an absolute CLR difference > 2.0 and were selected in ≥70% of bootstrap iterations.

For each significant pathway, constituent KOs were identified through a hierarchical mapping approach. MetaCyc pathway definitions were first decomposed into their constituent reactions and associated EC numbers, which were then mapped to KOs using the curated KEGG REST API. EC-level differential abundance was assessed using ALDEx2 (prevalence ≥ 10%, applying the same significance criteria as described above with an absolute CLR difference > 1.0). Only differentially abundant ECs were retained for downstream KO mapping, ensuring functional consistency between pathway- and gene-level signals. KO specificity was further refined by retaining only those exhibiting a Spearman correlation ≥ 0.3 with the corresponding pathway abundance across samples. KO-level differential abundance was then assessed using ALDEx2 with relaxed thresholds (prevalence ≥ 10%, CLR difference > 1.0, effect size ≥ 1.0). Taxon-level contributions to each KO were derived from PICRUSt2 stratified table, normalised within each sample by total KO abundance and aggregated at the genus level.

Results were visualised using multi-level network and bubble plots linking pathways, ECs, KOs, and contributing taxa. Pathway importance was summarised using a composite score incorporating perturbation rate, proportion of significance features, directional coherence, mean effect size, and KO coverage relative to total database-defined pathway gene content.

## Results

### Sample origin and snake clades

Out of 70 samples, 18 (25.7%) originated from snakes belonging to the Henophidia clade, while 52 (74.3%) belonged to the *Caenophidia* clade. The snakes originated from five private facilities, including Corboz (n = 25), Hausammann (n = 15), Grob (n = 13), Oberson (n = 10), and Eschlikon (n = 7) (Fig. 1a). Overall, 29 snakes (41.4%) were female and 41 (58.6%) were male (Fig. 1b). Within the *Henophidia* clade, nine snakes belonged to the family *Boidae* and nine to the family *Pythonidae*. Amongst the *Caenophidia* snakes, 45 individuals belonged to the family Viperidae, four to the Colubridae, and three to the Elapidae (Table S2).

**Fig. 1.**
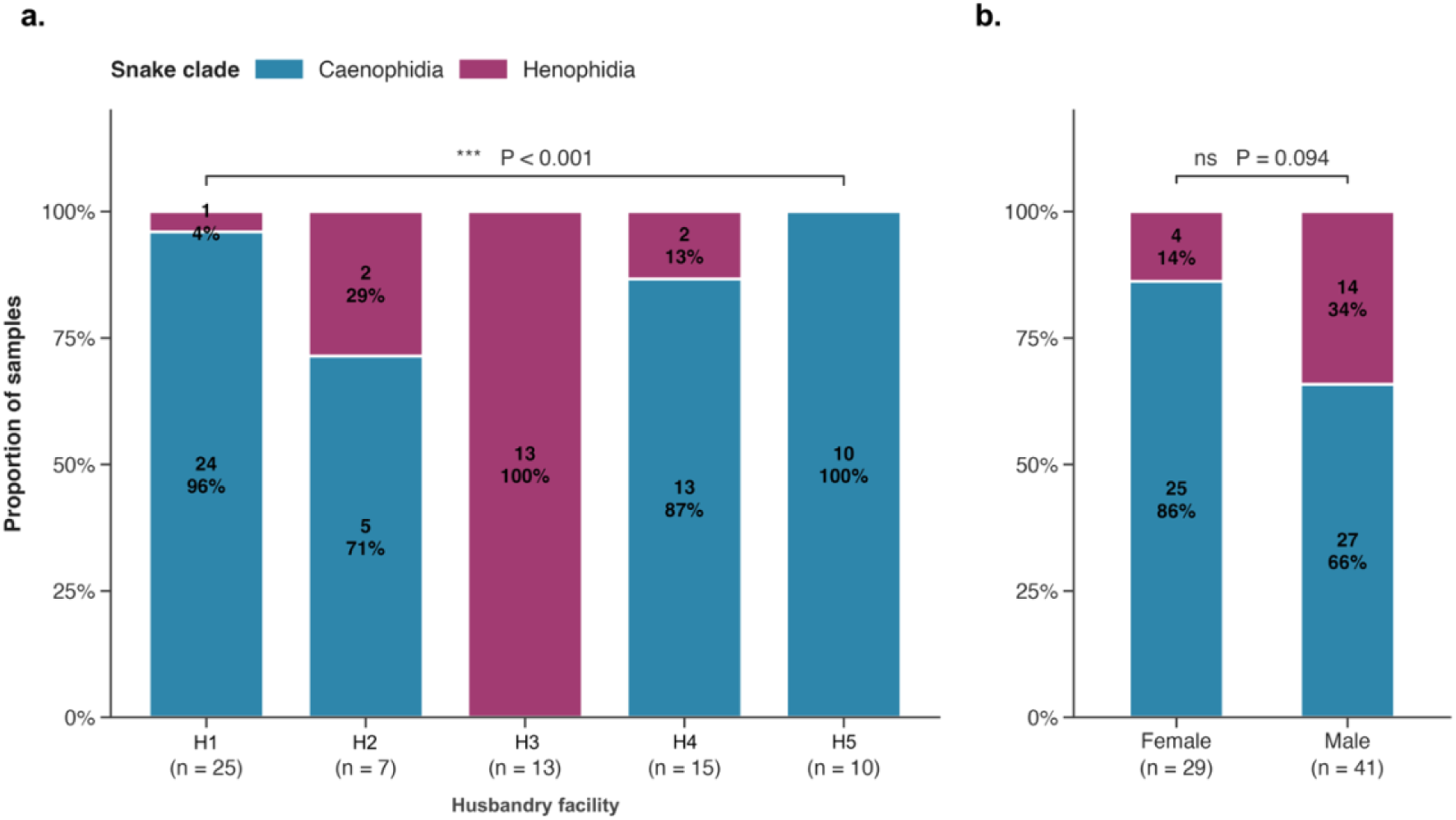
Distribution of snake clades across husbandry facilities and sex. Stacked bars show the proportional composition of *Caenophidia* and *Henophidia* within each stratum, with absolute sample counts and within-stratum percentages displayed inside each segment and the total number of samples per group reported below the corresponding axis label (n). (a) Clade composition across the husbandry facilities included in the study. (b) Clade composition between female and male animals. Statistical significance was assessed using Fisher’s exact test, testing the null hypothesis that clade membership is independent of husbandry facility (a) or sex (b). A significant result indicates that the relative proportions of *Caenophidia* and *Henophidia* differ across the categories shown.

### Henophidia display greater bacterial richness than Caenophidia

A total of 1967 ASVs were detected across all samples, with a mean of 70.0 ± 47.1 observed ASVs per sample (range: 17–249) (Table S1). Alpha diversity was assessed using three metrics; Chao1, Shannon and Pielou’s evenness. After controlling for confounding variables, *Caenophidia* exhibited significantly lower bacterial richness than *Henophidia* (Chao1: p = 0.006), with estimated marginal means of 78.3 (95% CI: 56.5-100.1) and 154.2 (95% CI: 114.7-193.7), respectively (Fig. 2a). No significant differences were observed between clades for Shannon diversity (p = 0.083) or Pielou’s evenness (p = 0.635) (Fig. 2b, 2c).

**Fig. 2.**
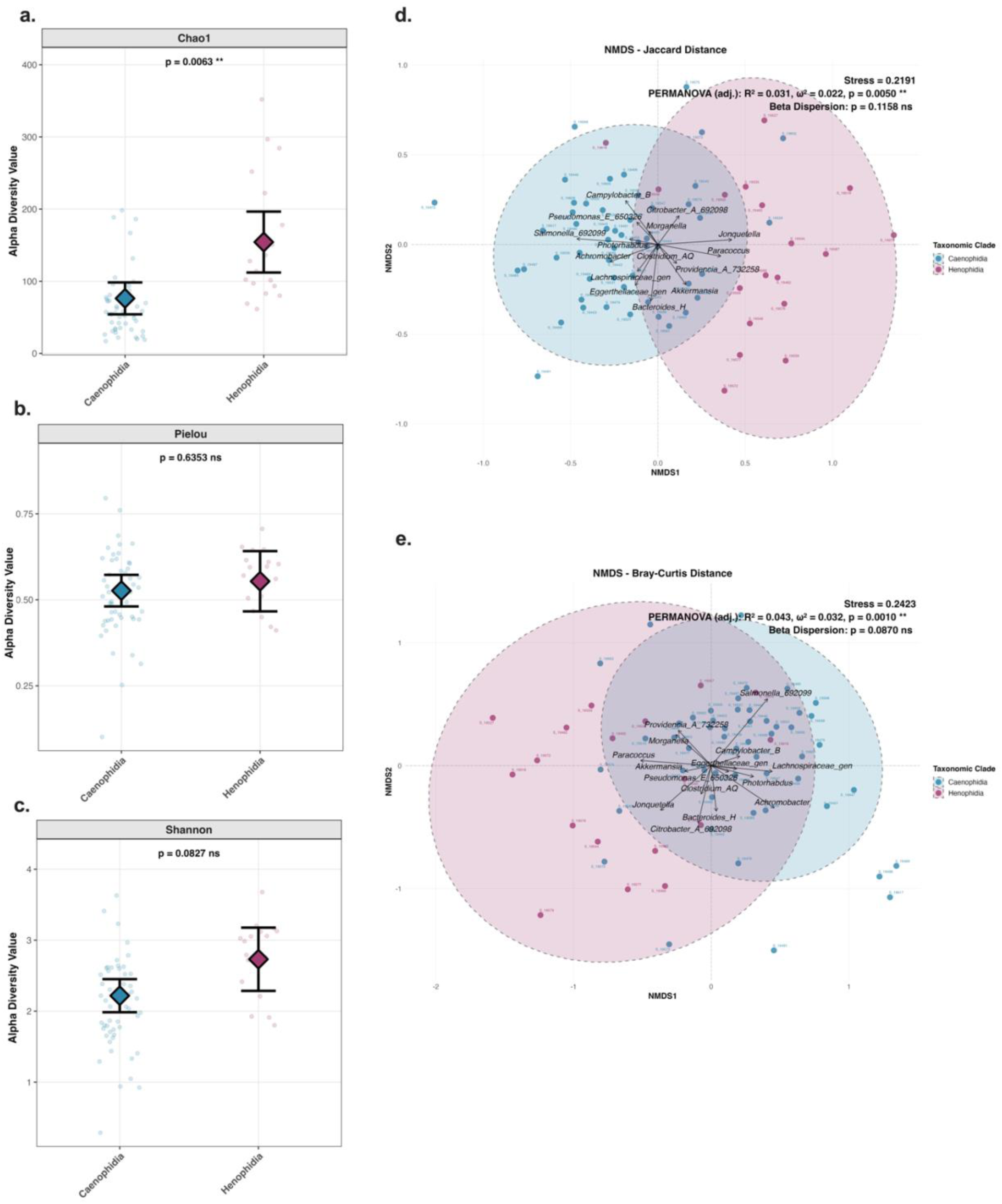
Microbial diversity comparisons between *Caenophidia* and *Henophidia* clades. (a-c) Estimated marginal means (±95% confidence intervals) for three alpha diversity metrics (calculated at the ASV level) including Chao1 bacterial richness (a), Pielou’s evenness (b), and Shannon diversity (c), adjusted for host sex and husbandry facilities. Large diamonds represent estimated marginal means, error bars indicate 95% confidence intervals, and individual sample values are shown as semi-transparent points. (d–e) NMDS ordinations at genus level using Jaccard (d) and Bray-Curtis (e) distance metrics. Each point represents an individual snake sample, coloured by clade (*Henophidia* = blue; *Caenophidia* = pink). Stress values quantify the goodness-of-fit between the original multidimensional distances and their two-dimensional representation. PERMANOVA R^2^ values represent the proportion of compositional variance explained by clade membership after accounting for sex and husbandry effects. Omega-squared (𝜔^2^) values indicate effect sizes accounting for unequal sample sizes. Genera vectors (black arrows) indicate the direction and magnitude of association between individual genera and the ordination space; arrow direction shows the gradient of increasing relative abundance for that genus, whilst arrow length reflects the strength of correlation with the ordination axes. Only genera with significant correlations to the ordination are displayed.

### Snake clade is a driver of cloacal microbiota composition

PERMANOVA testing on genus-level relative abundances demonstrated significant compositional differences between *Caenophidia* and *Henophidia* cloacal microbiota. Taxonomic clade explained 4.3% of the variance in Bray-Curtis (p=0.001, 𝜔^2^ = 0.032) and 3.1% in Jaccard (p=0.005, 𝜔^2^ = 0.022) distance matrices after controlling for sex and husbandry facilities (Fig. 2d and 2e, Table S3_1). Multivariate dispersion was homogeneous between clades for both Bray-Curtis (p=0.087) and Jaccard (p=0.116) distances, supporting the validity of PERMANOVA results for both metrics (Table S3_2). NMDS ordination revealed partial separation of clades along the first axis. Environmental vector fitting on Bray-Curtis ordination identified *Salmonella* (r^2^=0.46, p=0.001), *Achromobacter* (r^2^=0.33, p=0.001), *Jonquetella* (r^2^=0.27, p=0.001), *Paracoccus* (r^2^=0.26, p=0.001), *Citrobacter*_A (r^2^=0.23, p=0.001), *Lachnospiraceae*_gen (an unclassified genus within the family *Lachnospiraceae* for which Greengenes2 could not assign a formal genus-level name) (r^2^=0.20, p=0.004), *Providencia*_A (r^2^=0.14, p=0.005), *Bacteroides*_H (r^2^=0.14, p=0.006), *Morganella* (r^2^=0.13, p=0.005), and *Photorhabdus* (r^2^=0.10, p=0.032) as the taxa most strongly associated with clade separation along the ordination axes. For Jaccard distance, *Salmonella* (r^2^=0.22, p=0.003) and *Jonquetella* (r^2^=0.18, p=0.002) were the strongest contributors, with additional associations from *Paracoccus* (r^2^=0.13, p=0.018), *Bacteroides*_H (r^2^=0.10, p=0.029), and *Campylobacter*_B (r^2^=0.09, p=0.036) (Table S3_3).

### Husbandry facility, taxonomic clade, and sex partition cloacal community variation

To quantify how much of the variation in cloacal community composition was explained by taxonomic clade, sex, and husbandry facility, we used CAP with these variables as constraints. For Jaccard distance, the full model explained 17.0% of variance (adjusted R^2^=0.090, p=0.001) (Fig. 3a), with husbandry facility contributing 8.98% (p=0.001), clade 6.34% (p=0.001), and sex 1.85% (p=0.045) (Table S4_1). For Bray-Curtis distance, the full constrained model explained 15.0% of total variance (adjusted R^2^=0.070, p=0.001) (Fig. 3b), with marginal tests attributing 8.38% to husbandry facility (p=0.001), 4.23% to clade (p=0.001), and 2.31% to sex (p=0.014) (Table S4_2).

**Fig. 3.**
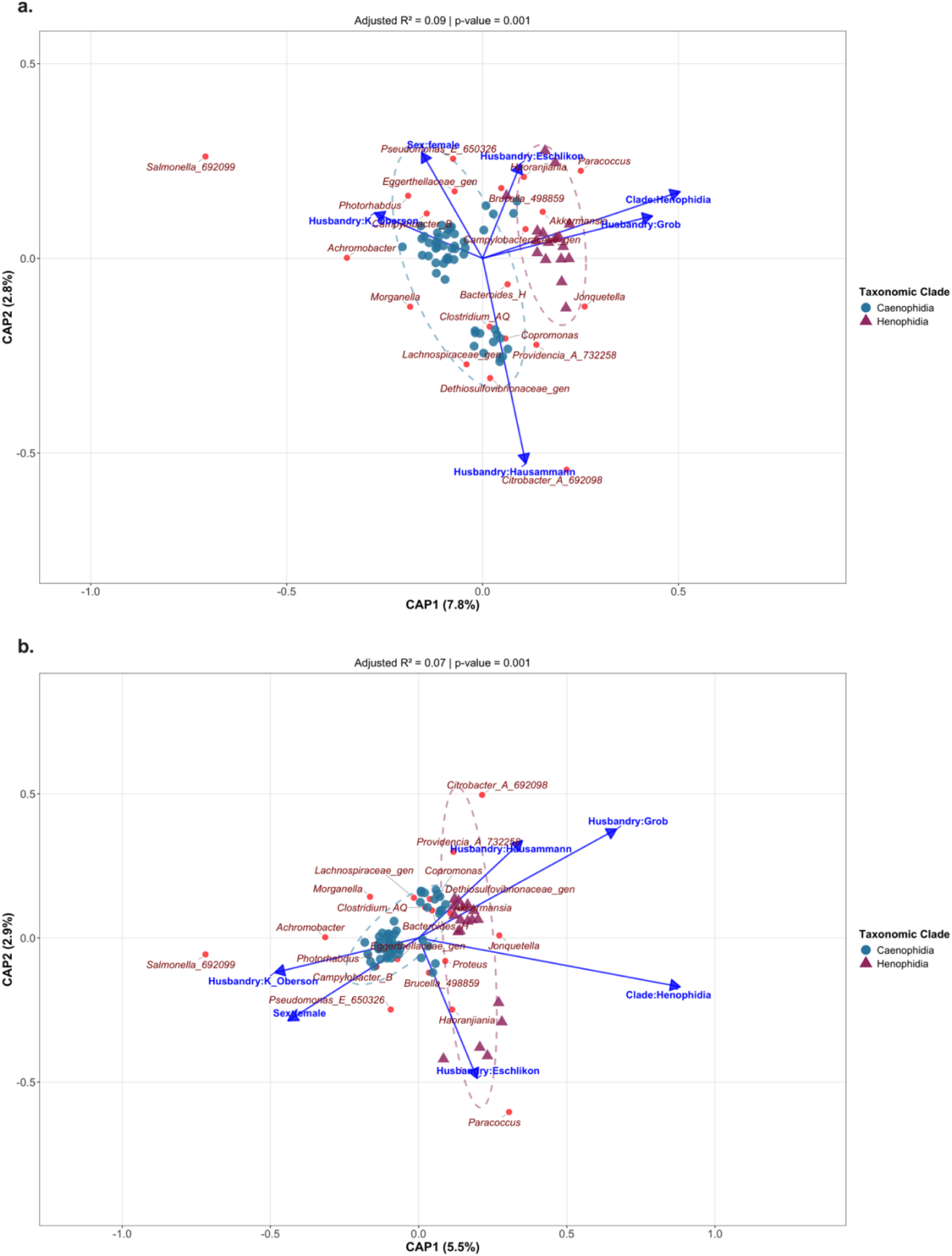
CAP ordination of cloacal microbiota composition based on (a) Bray-Curtis and (b) Jaccard distance. Each point represents an individual snake sample, coloured by taxonomic clade (*Caenophidia* = blue circles; *Henophidia* = pink triangles). Arrows represent fitted environmental variables (categorical predictor levels), with direction indicating the gradients of variation and length reflecting the strength of association with the ordination. Red points indicate the top 15 genera most strongly correlated with the ordination axes. Dashed ellipses enclose samples from each clade.

Post-hoc envfit identified genera whose abundance correlated with the constrained ordination axes, most strongly Salmonella under both Jaccard (r^2^=0.75, p=0.001) and Bray-Curtis (r^2^=0.72, p=0.001) distances, followed at lower r^2^ by *Citrobacter*_A, *Achromobacter*, *Paracoccus*, and *Jonquetella* (shared between both distances), with *Dethiosulfovibrionaceae*_gen and *Lachnospiraceae*_gen additionally significant under Jaccard, and *Providencia*_A and *Haoranjiania* under Bray-Curtis (Fig. 3a, 3b; Table S4_3, S4_4).

### Cloacal microbiota differs between snake clades across taxonomic levels

At the phylum level, both clades were dominated on average by *Proteobacteria*, followed by *Bacteroidota*, *Firmicutes*_A, and *Actinobacteriota* (Fig. 4a). At the genus level, 123 genera were detected across both clades after filtering, of which 37 were shared and encompassed the most abundant community members, whilst 3 were detected exclusively in *Caenophidia* and 83 exclusively in *Henophidia* (Fig. S1, S2; Tables S5_1-S5_3). *Salmonella* and *Bacteroides*_H were the most abundant genera overall, both showing higher mean relative abundance in *Caenophidia*, whilst genera such as *Morganella*, *Providencia*_A, and *Pseudomonas*_E were comparably abundant across clades (Fig. S1).

**Fig. 4.**
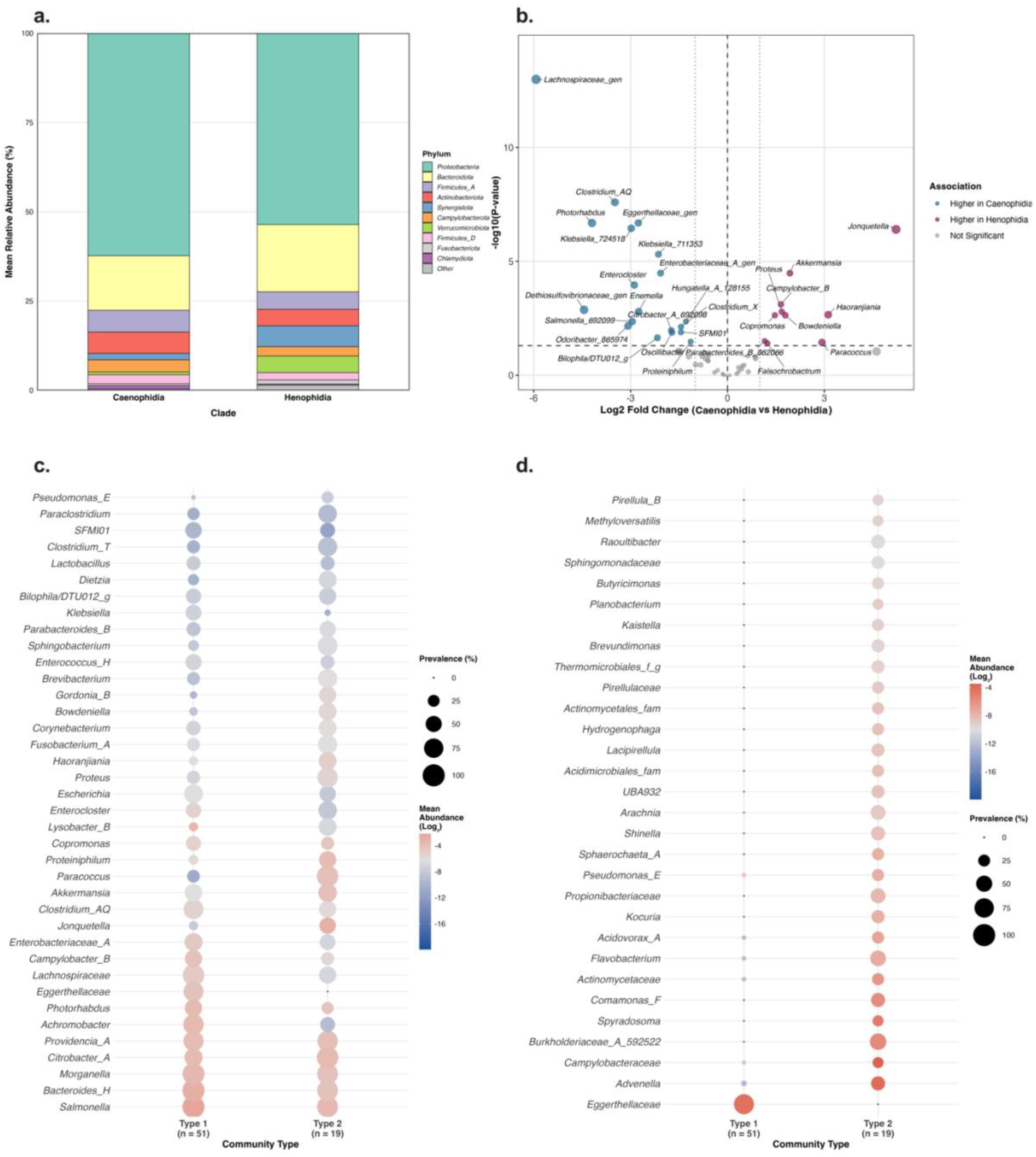
Taxonomic composition, differential abundance, and community-type structure of snake cloacal microbiota. (a) Phylum-level compositional profiles by clade. Stacked bar plots showing the mean relative abundance of the 10 most abundant phyla in *Caenophidia* and *Henophidia* cloacal microbiota. Each coloured segment represents a phylum, with height proportional to its mean relative abundance across all samples within that clade. Remaining phyla are grouped as “Other” (grey). (b) Differentially abundant genera between snake clades. Volcano plot showing differential abundance analysis results from ANCOM-BC2 comparing *Caenophidia* and *Henophidia* cloacal microbiota. Points represent individual genera coloured by association (blue: *Caenophidia*-enriched; pink: *Henophidia*-enriched; grey: not significant) based on FDR-adjusted q-values. Point size reflects effect size magnitude. Effect sizes represent log2FC in relative abundance after adjusting the linear model for sex and husbandry facility. The dashed horizontal line indicates the statistical significance threshold (q < 0.05); dotted vertical lines indicate the biological significance threshold (log2FC ≥ 1). (c, d) Driver (c) and discriminatory (d) taxa across DMM community types. (c) Dot plot of driver genera (n = 38), defined as those whose absolute fitted DMM parameter exceeded the 80^th^ percentile in at least one community type. (d) Dot plot of discriminatory genera (n = 30), selected as the union of the top genera per community type ranked by the absolute log2FC in mean relative abundance between that community type and all other samples. In panels (c) and (d), point size denotes prevalence (percentage of samples within each community type in which the taxon was detected) and colour denotes mean relative abundance on a log2 scale, calculated only from samples in which the taxon was present. Genera are ordered along the y-axis by their overall mean relative abundance across all samples in the dataset (zero values included).

Because mean relative abundances and presence/absence patterns are descriptive and may be confounded, we used ANCOM-BC2 to test for differential abundance between clades whilst adjusting for sex and husbandry facility. After adjustment, 29 genera differed significantly between clades (q<0.05), with the strongest opposing signals being *Lachnospiraceae*_gen (log2FC = -5.93, q = 1.0x10^-13^) higher in *Caenophidia*, alongside *Dethiosulfovibrionaceae*_gen and multiple *Enterobacteriaceae* (*Photorhabdus*, *Klebsiella*, *Salmonella*, *Citrobacter*_A), and *Jonquetella* (log2FC = +5.22, q = 3.9x10^-7^) higher in *Henophidia*, accompanied by *Haoranjiania*, *Paracoccus*, and *Akkermansia* (Fig. 4b; Table S6).

### Two cloacal bacterial community types align with snake clade

To assess whether snake cloacal microbiota partition into discrete community states independently of clade labels, we applied DMM modelling. The optimal solution comprised two community types (k=2; Laplace = 13,344.26): Type 1 (n=51) was almost entirely *Caenophidia* (49/51; 96.1%), whilst Type 2 (n=19) was predominantly *Henophidia* (16/19; 84.2%) (Fig. S3; Fig. S4), indicating that unsupervised community structure aligns closely with snake clade. The five samples that did not align with their taxonomic clade were inspected individually, but no consistent association with husbandry facility, host sex, or finer taxonomic resolution (family level) could be identified to explain the misalignment.

To characterise each community type, genera were summarised in two complementary ways: by their fitted Dirichlet parameters from the DMM model, which quantify each genus’s contribution to the expected composition of that community type (driver taxa; Fig. 4c) and by absolute log2FC in mean abundance between types, identifying the genera showing the largest between-type differences (Fig. 4d). Type 1 was driven primarily by *Salmonella*, *Bacteroides*_H, *Morganella*, *Lachnospiraceae*, and *Providencia*_A, whilst Type 2 was driven by *Paracoccus*, *Bacteroides*_H, *Morganella*, *Providencia*_A, and *Citrobacter*_A; several genera (*Bacteroides*_H, *Morganella*, *Providencia*_A) ranked as top drivers in both types at different contribution levels, whereas others were largely community-specific (*Salmonella* in Type 1, *Paracoccus* in Type 2). The fold-change ranking highlighted *Eggerthellaceae* as the genus most strongly associated with Type 1, and *Burkholderiaceae*_A, *Spyradosoma*, *Comamonas*_F, and *Advenella* as those most associated with Type 2. Consistent with the community types reflecting clade structure, the top drivers of each type substantially overlapped with the genera identified as clade-discriminating by ANCOM-BC2, including *Lachnospiraceae*, *Eggerthellaceae*, *Photorhabdus*, *Klebsiella*, and *Clostridium*_AQ in Type 1, and *Paracoccus*, *Akkermansia*, *Haoranjiania*, *Jonquetella*, and *Bowdeniella* in Type 2.

### Henophidia microbiota show denser, more modular co-occurrence networks

To examine whether genera co-occur differently between snake clades, we constructed clade-specific co-occurrence networks (SparCC) at the genus level. The two networks differed in connectivity and organisation (Fig. 5; Table S7_1). The *Caenophidia* network was sparse, with very low edge density (0.0007) and exclusively positive associations, whereas the *Henophidia* network was 11.9-fold denser (0.0083) and remained predominantly positive (93.5%). The two networks also differed in topology: *Henophidia* showed higher modularity (0.80 *vs*. 0.32), indicating clearer partitioning into sub-communities, whereas *Caenophidia*’s higher clustering coefficient (0.78 *vs*. 0.49) reflects only the few connected genera within its otherwise sparse network.

**Fig. 5.**
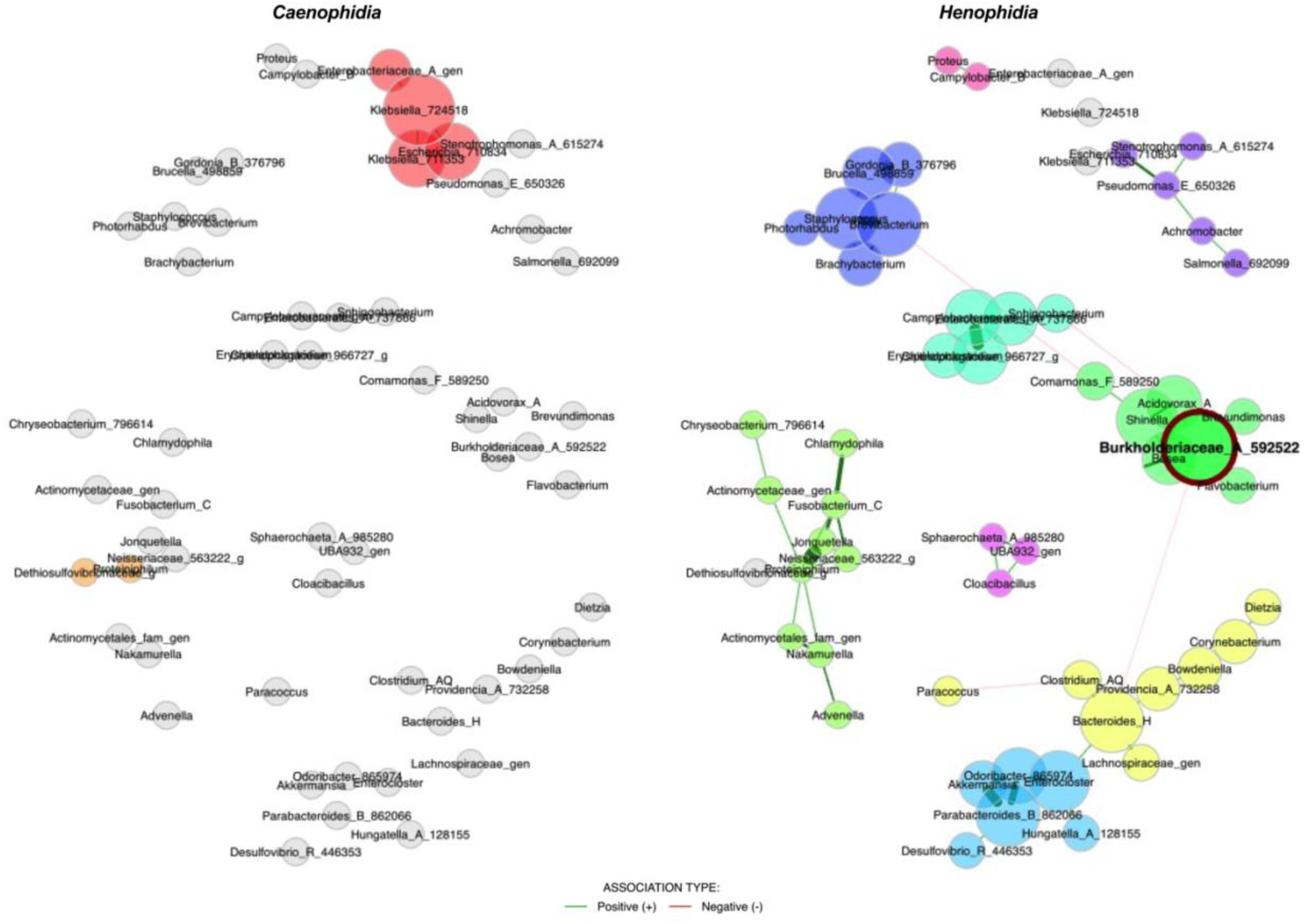
Microbial co-occurrence network in *Caenophidia* and *Henophidia*. SparCC-inferred genus-level networks for *Caenophidia* and *Henophidia*. Nodes represent genera sized by degree centrality. Module detection was performed independently within each network using fast greedy modularity optimisation; node colours therefore indicate within-network modules and do not correspond between panels. Hub genera are outlined in red. Edges represent strong, statistically significant associations (r ≥ 0.5, lfdr < 0.05), with edge thickness proportional to the magnitude of the correlation coefficient (thicker edges indicating stronger co-occurrence patterns) and colour indicating association type.

*Caenophidia* network were largely unstructured, with most genera (117/123) grouped into a single weakly structured cluster and only two small modules, a four-member *Enterobacteriaceae* cluster and a two-member cluster containing *Proteiniphilum* and *Dethiosulfovibrionaceae* (Fig. 5; Table S7_2). In contrast, *Henophidia* network resolved into ten distinct modules, including an anaerobe-enriched cluster (*Bacteroides*, *Bowdeniella*, *Clostridium*, *Lachnospiraceae*) and a *Burkholderiaceae*-centred module, alongside several smaller specialised clusters of 2–10 genera each (Fig. 5; Table S7_3). Henophidia network also displayed greater nodal connectivity, with an unclassified *Burkholderiaceae* genus identified as the most central hub (degree = 2.78, betweenness = 0.58) and additional highly connected taxa (e.g., *Brevibacterium*, *Parabacteroides*, *Bacteroides*_H, *Staphylococcus*) listed in Table S7_3. Pairwise Jaccard similarity between *Caenophidia* and *Henophidia* modules was uniformly low (maximum = 0.13), with only *Escherichia* and *Proteiniphilum* appearing in any module common to both networks, indicating that the two networks differ not only in topology but also in modular composition.

### Inferred functional differences between clades are subtle but specific

To assess whether the two clades differed in their predicted functional potential, we used PICRUSt2 to infer MetaCyc pathway abundances from the 16S rRNA gene profiles. PERMANOVA on Bray-Curtis distance showed that clade explained a non-significant fraction of the variance in predicted functional composition (1.9%; p=0.196, 𝜔^2^=0.002) and betadisper indicated no significant differences in multivariate dispersion for any variable (p>0.16; Table S8_1).

Despite the absence of global functional separation, two complementary analyses were applied to identify pathways differing between clades, each addressing a distinct question. ALDEx2 differential abundance testing identified 23 pathways significantly differing between clades, of which 15 were more abundant in *Henophidia* and eight in *Caenophidia* (Fig. 6; Table S8_2). Bootstrap-stabilised elastic net regression, controlling for husbandry facility and sex as unpenalised covariates, selected 24 features that were stably informative for distinguishing the two clades across 100 bootstrap iterations; of these, five met the CLR difference > 2.0 threshold, all with higher CLR abundance in *Henophidia* (Fig. 6; Table S8_3, S8_4). Four pathways (ethylmalonyl-CoA pathway, formaldehyde assimilation I, glycine betaine degradation I, and methylaspartate cycle) were consistently identified across both differential abundance testing and feature selection (Fig. 6).

**Fig. 6.**
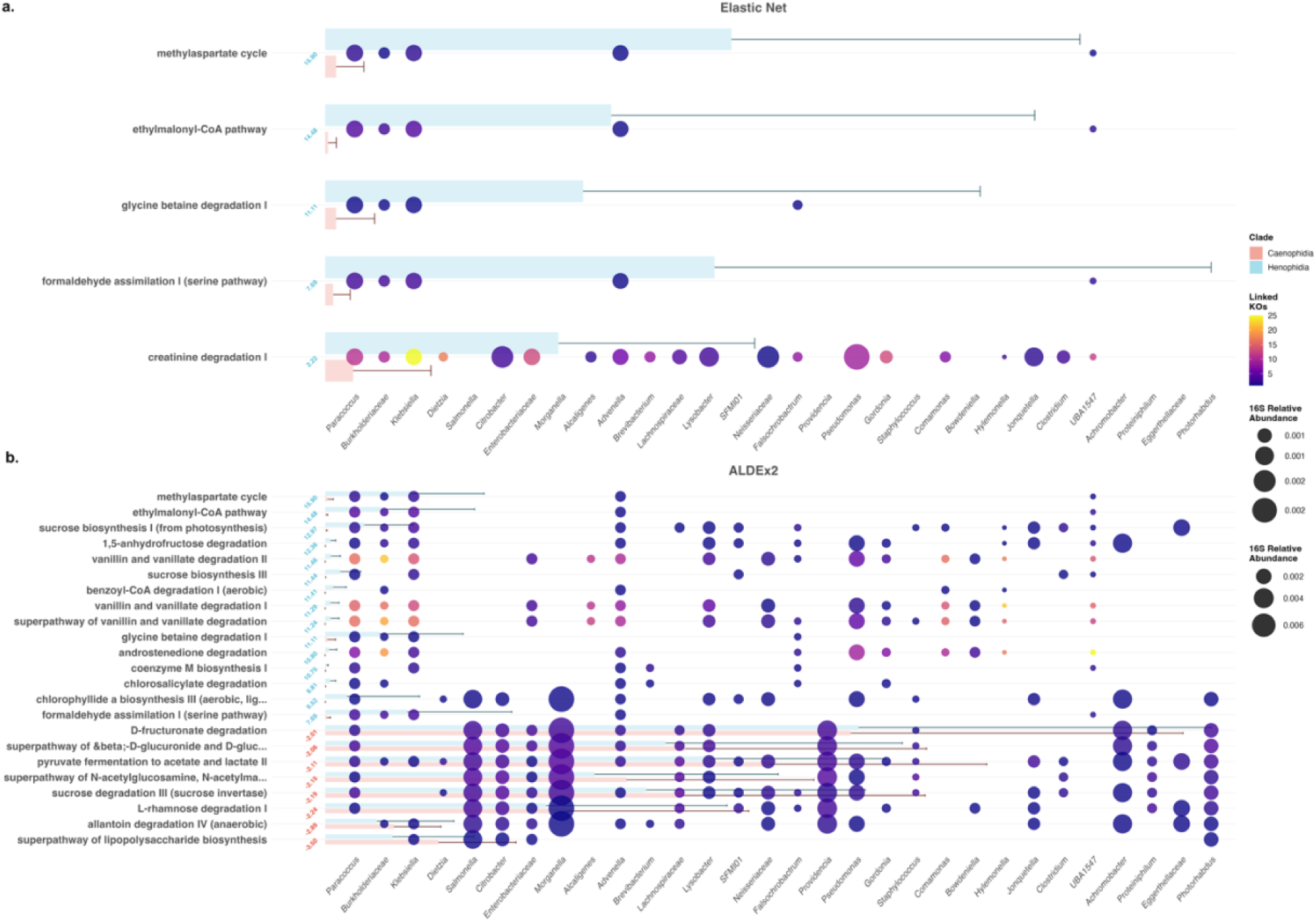
Differentially abundant MetaCyc pathways and their taxonomic contributions; (a) Elastic net–selected and (b) ALDEx2-identified. Pathways are ordered by CLR difference between *Caenophidia* and *Henophidia*, with CLR difference values displayed at 45° adjacent to the y-axis and coloured by enrichment direction. Transparent horizontal bars represent mean predicted pathway abundance (± SD) per clade. Genera (x-axis, italic) represent the top 30 taxa contributing to the displayed pathways via KEGG Ortholog (KO)-mediated linkage, ordered by total abundance. Bubbles at each pathway-taxon intersection indicate a functional connection mediated by one or more KOs; bubble size is proportional to the mean 16S rRNA gene-derived relative abundance of that genus across all samples, whilst bubble colour intensity reflects the number of distinct KOs linking each taxon to the pathway, with brighter colours denoting a broader enzymatic contribution. Only pathways with ≥ 60% prevalence in at least one clade and meeting method-specific significance thresholds (ALDEx2: adjusted p < 0.05, effect size ≥ 1.0, CLR difference > 2.0; Elastic Net: CLR difference > 2.0, selection frequency ≥ 70% across 100 bootstrap iterations) are shown.

To explore the taxonomic contributors underlying functional differences, pathways identified as differentially abundant by ALDEx2 and pathways selected by the elastic net model as predictive features were decomposed into their constituent KOs and linked to contributing genera (Table S8_5–S8_6). Linkage was successfully established for all elastic net (5 pathways; 31 KOs; 510 linkages) and all ALDEx2 pathways (23 pathways; 98 KOs; 1,983 linkages), confirming complete pathway-KO-taxon traceability across both methods.

Across both methods, pathways more abundant in *Henophidia* were supported by contributions from a broader set of taxa and KOs, whereas pathways more abundant in *Caenophidia* showed more concentrated contributions from fewer taxa (Fig. 6; Tables S8_5-S8_6). *Henophidia*-associated pathways also showed substantially larger ALDEx2 effect sizes than *Caenophidia*-associated pathways (mean 11.45 *vs*. 2.14). Examples of this pattern included the ethylmalonyl-CoA pathway and formaldehyde assimilation I, both more abundant in *Henophidia* and supported by multiple contributing genera, whereas the superpathway of β-D-glucuronide and D-glucuronate degradation was more abundant in *Caenophidia* (Fig. S5). Among the elastic net features, creatinine degradation I emerged as the most extensively connected pathway, with contributions concentrated in *Henophidia*-associated taxa.

At the genus level, a small number of taxa contributed disproportionately to pathway-level differences, including members of *Burkholderiaceae*, *Paracoccus*, and *Candidatus* Saccharibacteria UBA1547. Several genera showed clade-specific contributions: *Candidatus* Saccharibacteria UBA1547, *Hylemonella*, and *Comamonas* contributed almost exclusively to *Henophidia*-associated linkages, whereas *Klebsiella* contributed predominantly to *Caenophidia*-associated pathways.

### Integration of community structure, networks, and functional profiles

To synthesise across analyses, we focused on genera that emerged consistently across multiple analytical layers, clade-level differential abundance, DMM driver and discriminatory taxa, network centrality and module membership, and pathway–taxon linkage analysis, to identify taxa with convergent multi-method support. *Henophidia* (Type 2) communities were characterised by broad, coordinated functional contributions distributed across many taxa, whereas *Caenophidia* (Type 1) communities showed narrower contributions concentrated in fewer taxa. In *Henophidia*, the unclassified *Burkholderiaceae* genus served as the network hub and, together with *Paracoccus*, ranked as top driver and DMM discriminatory taxa whilst also being the most broadly connected taxa in the pathway–taxon linkage analysis, contributing across a wide range of metabolic functions including aromatic compound degradation. Additional Henophidia-associated taxa, including *Comamonas*, *Advenella*, *Candidatus* Saccharibacteria UBA1547, and *Hylemonella*, appeared exclusively in *Henophidia*-associated functional linkages, reinforcing the clade specificity of this functional signature.

In contrast, the *Caenophidia* functional signature was concentrated in fewer taxa, with *Klebsiella,* the only network-connected genus among the top *Caenophidia* drivers, dominating contributions to a small number of pathways. *Lachnospiraceae* and *Eggerthellaceae*, the strongest ANCOM-BC2 signals on the *Caenophidia* side, showed more limited pathway involvement. Several genera (*Morganella*, *Providencia*_A, *Salmonella*, *Citrobacter*_A) ranked as drivers in both DMM types and contributed comparably across clades, consistent with a conserved core of community members supporting shared metabolic functions whilst clade-specific taxa account for the functional divergence between clades.

## Discussion

This study characterised the cloacal microbiota of 70 captive snakes, suggesting that *Caenophidia* and *Henophidia* harbour compositionally and functionally distinct microbial communities despite sharing a core set of dominant genera. *Caenophidia* exhibited substantially lower bacterial richness than *Henophidia*, whilst Shannon diversity and Pielou’s evenness remained comparable, suggesting that the two clades differ in the number of taxa detected but may share a similar dominance structure and overall evenness [34]. This pattern is consistent with prior findings: Zhang *et al*. [35] demonstrated interspecific variation in microbial richness amongst farmed snakes, and Hoffbeck *et al*. [36] reported the taxonomic order as one of the strongest contributors to alpha diversity variation across 91 reptile species. Regarding beta diversity, we recorded compositional differences between clades for both Bray-Curtis and Jaccard distance matrices. This clade-associated variation is broadly consistent with other reptile systems: Dallas *et al*. [37] identified host taxonomy as the primary determinant of cloacal microbiota composition in nearctic colubrids, Zhang *et al*. [35] showed that conspecific microbiotas in snakes clustered together despite shared captive conditions, and Smith *et al*. [38] reported compositional separation amongst venomous snake species driven by both ecological and phylogenetic factors. The partial NMDS separation of *Caenophidia* and *Henophidia*, together with the identification of *Bacteroides_H*, *Salmonella*, *Achromobacter*, and *Jonquetella* as the genera most strongly correlated with the ordination axes, suggests that phylogenetic clade membership may structure cloacal microbiota composition whilst leaving substantial residual variance attributable to other factors.

The cloacal microbiota of both clades was dominated at the phylum level by *Proteobacteria*, followed by *Bacteroidota*, *Firmicutes_*A, and *Actinobacteriota*, a compositional profile consistent with the established core microbiome of snakes reported across diverse species [7,23,35,39–41], and confirmed by a meta-analysis of 91 reptile species [36]. At the genus level, *Salmonella* was the most abundant taxon overall yet displayed pronounced clade asymmetry (20.3% in *Caenophidia vs*. 8.6% in *Henophidia*), consistent with its widespread detection across captive and farmed snake species [8,35,40,42]. Differential abundance testing with ANCOM-BC2 results align with these compositional patterns, identifying 29 clade-associated genera, with *Caenophidia* enriched in multiple *Enterobacteriaceae* members (*Salmonella*, *Klebsiella*, *Photorhabdus*), and *Henophidia* enriched in *Akkermansia*, as well as *Jonquetella* and *Paracoccus*. The markedly higher number of clade-specific genera in *Henophidia* compared with *Caenophidia* aligns with the greater bacterial richness observed in the former clade and suggests that boid and pythonid hosts may support a broader array of microbial lineages [7,21].

DMM modelling identified two community types that closely corresponded to the phylogenetic clades, although the concordance was not complete. This incomplete correspondence may reflect the influence of confounding factors, particularly husbandry facility and host sex. In captive settings, such environmental heterogeneity has been shown to influence gut microbiota across vertebrates, with captivity-associated changes driven largely by case-specific conditions rather than universal patterns [43]. Previously, Dallas *et al*. [37] reported that host taxonomy was the primary driver of cloacal microbiome composition in colubrids. Driver taxa analysis revealed that the distinction between community types was driven by shifts in the relative dominance of shared genera, such as *Bacteroides*_H, *Morganella*, and *Providencia*_A, which were present in both types but at different abundance levels. Type 1 was further characterised by *Salmonella* and showed the largest difference in *Eggerthellaceae* abundance, whilst Type 2 was characterised by *Paracoccus* and the largest differences in *Burkholderiaceae*_A, *Comamonas*_F, and *Advenella*. The Type 2-associated genera, *Burkholderiaceae*_A, *Comamonas*_F, *Advenella*, were not reported in the prior studies on henophidian snakes [7,24]. The cloacal microbiota of boid and pythonid hosts therefore remain sparsely characterised, and we cannot yet determine whether these genera reflect features genuinely associated with henophidian snakes or features specific to this cohort.

A central question arising from our findings is the relative contribution of host phylogeny *vs*. environmental factors to the observed microbiota differences. CAP analysis quantified this by partitioning compositional variance among predictors: for both Bray–Curtis and Jaccard distances, husbandry facility consistently explained more variance than clade or sex, suggesting that husbandry facility exerts a stronger influence on cloacal microbiota than the phylogenetic divergence between *Henophidia* and *Caenophidia*. This might be consistent with the few cross-classified individuals in the DMM analysis, in whom environmental factors could have partially modulated, rather than fully overridden, the phylogenetic signal. Nonetheless, the clade effect remains substantial, and its true magnitude may be underestimated owing to confounding between clade and husbandry facility. Our findings sit within an active debate over the relative importance of phylogeny, diet and environment in shaping reptilian gut microbiota: Hoffbeck *et al*. [36] identified host diet and taxonomic order as the strongest predictors across 91 reptile species, Youngblut *et al*. [21] demonstrated that phylogeny and diet modulate different aspects of vertebrate gut microbiome diversity, and Song *et al*. [20] and Zhu *et al*. [9] showed that dietary specialisation may exceed phylogenetic relatedness in wild snakes, observations broadly in line with our findings. The captivity context further moderates this balance: Huang *et al*. [44] reported that captivity diminishes the effects of host phylogeny on gut bacterial diversity across vertebrates, and Xie *et al*. [45], in a meta-analysis of 113 vertebrate species, identified diet as a major driver of microbiome variation in wild animals, noting that diet and phylogeny are not independent because evolutionary history shapes the dietary niches available to host lineages. By contrast, Zhang *et al*. [35] reported that phylogenetic signal can persist even under standardised husbandry in farmed snakes, suggesting that the balance between environmental and phylogenetic effects depends on species, husbandry conditions, and the microbiome compartment under study.

Co-occurrence network analysis revealed notable differences between the two clades. The *Caenophidia* network was remarkably sparse, with only six genera forming significant co-occurrence relationships, however, these formed exclusively positive, tight associations, reflected in a high clustering coefficient (0.778), suggesting that co-occurring taxa in this clade interact with strong specificity despite overall low connectivity. By contrast, the *Henophidia* network was 11.9-fold denser, being resolved into 10 distinct modules with more balanced size distributions, and featured a *Burkholderiaceae* hub genus, with a longer average path length (2.459 *vs.* 0.603) consistent with its distributed modular topology. Higher network modularity, as observed in *Henophidia*, is generally interpreted as indicative of a community partitioned into sub-communities of taxa that interact more strongly with one another than with taxa in other modules [46,47]. Such modular architecture is thought to confer greater community stability, as perturbations affecting one module are less likely to propagate to the rest of the network [48,49]. The functional interpretation of these modules as ecological guilds is supported by literature on their member taxa: the *Burkholderiaceae*-centred module groups genera from a family well-characterised for aerobic aromatic-substrate catabolism via the benzoate oxidation cluster [50–52], whilst the anaerobe module (*Bacteroides*, *Bowdeniella*, *Clostridium*, *Lachnospiraceae*) consists of genera widely documented as obligate fermentative anaerobes of the vertebrate gut [53,54]. The sparse *Caenophidia* network may reflect a community assembled through stochastic or environmental constraints. Amongst the limited snake microbiome network studies available, Zhu *et al*. [9] constructed co-occurrence networks for three colubrid species with distinct diets. They found that the invertebrate-feeding species exhibited the most complex network with the highest number of nodes, links, and negative associations, whilst the two vertebrate-feeding species displayed more stable networks with higher modularity and lower competitive interactions. This diet-dependent variation in network complexity may in part be in line with our clade-level findings, as *Henophidia*, which consume larger, less frequent meals, harboured a denser and a more modular network, potentially reflecting the greater ecological complexity required to sustain microbial communities through prolonged fasting-feeding cycles.

The predicted functional differences between clades, although modest, revealed specific pathway-level enrichments that aligned in part with the differential abundance and network analyses to identify a coherent set of taxa simultaneously driving compositional differentiation, occupying structurally central network positions, and revealing clade-specific metabolic capacity. The larger ALDEx2 effect sizes associated with *Henophidia* pathways than *Caenophidia* may suggest that functional distinctions were driven by strong, directional *Henophidia*-specific enrichments rather than bidirectional differentiation of comparable magnitude. The four pathways enriched through both differential abundance testing and feature selection in *Henophidia*, ethylmalonyl-CoA pathway, formaldehyde assimilation I, glycine betaine degradation I, and methylaspartate cycle, suggest enhanced C1 metabolism and alternative carbon assimilation strategies relevant to the distinctive digestive physiology of pythons and boas, which undergo postprandial organ remodelling followed by prolonged fasting during which the gut atrophies [7,19]. These C1 pathways may support microbial survival during nutrient scarcity, enabling utilisation of host-derived substrates such as glycine betaine from mucosal turnover. By contrast, *Caenophidia*-enriched pathways, including allantoin degradation IV and the superpathway of β-D-glucuronide and D-glucuronate degradation, suggest greater microbial capacity for purine catabolism and carbohydrate processing, potentially reflecting the more frequent feeding patterns of viperids and colubrids. Two further *Henophidia*-enriched pathways identified in our analysis, chlorosalicylate degradation and benzoyl-CoA degradation I (aerobic), reinforce a broader capacity for aerobic aromatic compound catabolism in this clade. Benzoyl-CoA degradation is the central aerobic route for benzoate and halogenated aromatic substrates in bacteria and has been characterised extensively in members of the *Burkholderiales*, where it involves the box (benzoate oxidation) enzymatic cluster [50,51]. The *Burkholderiaceae*-centred network module and the anaerobe-enriched module (*Bacteroides*, *Bowdeniella*, *Clostridium*, *Lachnospiraceae*) may represent functionally distinct ecological guilds involved in aromatic compound degradation and fermentative processes, respectively, consistent with the vanillin degradation and C1 pathway enrichments. Creatinine degradation I was the most extensively connected pathway, with a pronounced *Henophidia* bias in contributing taxa. Creatinine is a nitrogenous waste product whose microbial degradation constitutes an important mechanism of intestinal nitrogen recycling [55], and the *Henophidia*-biased contributor community is consistent with the greater physiological relevance of nitrogen conservation during the prolonged fasting periods characteristic of pythons and boas [56,57].

*Burkholderiaceae* exemplifies the convergence amongst different analysis approaches: (*i*) it was significantly enriched in *Henophidia*, (*ii*) served as the sole hub genus in the *Henophidia* co-occurrence network, (*iii*) was the most discriminatory taxon for DMM Type 2 communities, and (*iv*) emerged as the most broadly connected genus across differentially abundant pathways, suggesting it functions as a keystone taxon in the *Henophidia* cloacal microbiome [58]. *Akkermansia* is a specialised mucin-degrading bacterium, whose enrichment in *Henophidia*, likely reflects prolonged fasting periods, consistent with its expansion in fasting Burmese pythons [7] and the isolation of *Akkermansia glycaniphila* from the reticulated python gut [59,60]. *Jonquetella*, nearly exclusive to *Henophidia*, is an obligate anaerobe associated with amino acid fermentation relevant to the prolonged gut transit times of large-bodied constrictors. By contrast, the enrichment of *Enterobacteriaceae* members (*Salmonella*, *Klebsiella*, *Photorhabdus*) in *Caenophidia*, with *Klebsiella* contributing predominantly to *Caenophidia* functional linkages, may reflect the higher feeding frequency of viperids and colubrids, favouring fast-growing, facultatively anaerobic taxa. This biological coherence, mucin-degrading and amino acid-fermenting taxa suited to prolonged fasting in *Henophidia*, and *Enterobacteriaceae* suited to frequent feeding in *Caenophidia*, together with the concordance across compositional, network, and functional analytical layers, strengthens the inference that clade differences may reflect an interconnected system in which taxonomic identity, microbial interactions structure, and metabolic capacities are jointly organised along phylogenetic lines. Certain taxa showed discordant patterns: *Eggerthellaceae* was the most discriminatory taxon for Type 1 communities yet displayed limited functional connectivity, highlighting a recognised limitation of PICRUSt2-based inference for some poorly characterised taxa [61].

Several limitations should be considered for this study. First, the unequal sample sizes between *Caenophidia* and *Henophidia* may have reduced the statistical power to detect subtle compositional and functional differences. This imbalance also meant that within-clade diversity estimates for *Henophidia* were based on fewer observations, potentially inflating confidence intervals and limiting the detection of rare taxa. Second, all functional predictions were derived from PICRUSt2, which infers metabolic potential from 16S rRNA gene data. The functional differences reported here should be regarded as hypothesis-generating, and validation through shotgun metagenomic sequencing is recommended. Third, the association between husbandry facility and clade composition means that residual confounding between these variables cannot be fully excluded. Fourth, the captive origin of all sampled snakes limits the generalisability of our findings to wild populations. Captivity has been shown to alter gut microbiota composition across diverse vertebrate taxa, often through homogenisation of microbial communities and enrichment of human-associated microorganisms [43,44].

Integrating compositional, network, and predicted-functional analyses across 70 captive snakes reveals coherent microbiota differences between *Caenophidia* and *Henophidia* that persist alongside, though partly subordinate to, the influence of husbandry facility. These results indicate that the cloacal microbiota of captive snakes is probably structured in part by phylogenetic clade membership. *Burkholderiaceae* emerges as a candidate keystone taxon in *Henophidia*, supported by convergent signals across all analytical layers. The lower bacterial richness observed in *Caenophidia* raises a testable hypothesis: that the broad-spectrum antimicrobial activity of caenophidian venom components, including phospholipase A2 enzymes, L-amino acid oxidases, and cationic peptides [62], may impose a selective filter on susceptible microbial lineages. Whether the microbiota differences observed here reflect long-standing host-microbe coevolution [63] or are more proximately shaped by clade-associated differences in digestive physiology and feeding ecology cannot be resolved with the present data. Future work incorporating wild-caught individuals, controlled dietary experiments, longitudinal sampling across feeding-fasting cycles, and shotgun metagenomic approaches will be needed to disentangle these contributions. By extending multi-method microbiome characterisation to underrepresented henophidian hosts, this study provides an integrated baseline for a broader research programme on the interplay between host phylogeny, ecology, and microbial community structure across snake diversity.

## Resource availability

### Lead contact

Professor Gilbert Greub.

### Materials availability

All materials will be made available upon request.

### Data and code availability

16S rRNA gene amplicon sequencing data generated in this study will be deposited at the European Nucleotide Archive prior to publication. Accession numbers will be provided upon acceptance and will be publicly available as of the date of publication. All original code used for sequence processing, statistical analysis, and figure generation will be deposited at Zenodo prior to publication, and the DOI will be provided upon acceptance. Any additional information required to reanalyse the data reported in this paper is available from the lead contact upon request.

### Ethics approval and consent to participate

The collection and molecular analysis of the snake samples was approved and performed in accordance with the relevant guidelines and regulations of the Veterinary Office of Canton Zurich (authorization no. ZH010/15).

### Consent for publication

All authors consent to the publication of this manuscript.

### Author contribution

E.G. manuscript drafting and data analysis; S.N. data curation and methodology; T.P. data acquisition; S.R. data acquisition; S.A. data acquisition; C.B. design of the work; N.B. design of the work and funding acquisition; G.G. design of the work and funding acquisition; All authors reviewed the manuscript.

### Funding

This work was partially funded by the SNSF-NCCR microbiome grant to Gilbert Greub (IMUR 28881). This work was also partially supported by the Swiss Association for Wildlife, Zoo and Exotic Animals to NB.

## Acknowledgments

We gratefully acknowledge the sequencing facility of the DMLP at Lausanne University Hospital for their technical support and 16S rRNA amplicon sequencing, and Ms Rizlène Dira for her assistance with sample preparation. We extend our thanks to Dr Niklaus Johner and Elindi de Coning for insightful discussions.

