## Supplementary Data for "Comparative analysis of cloacal microbiota in *Henophidia* (non-venomous) and *Caenophidia* (venomous) snakes"

**Category**

Original article

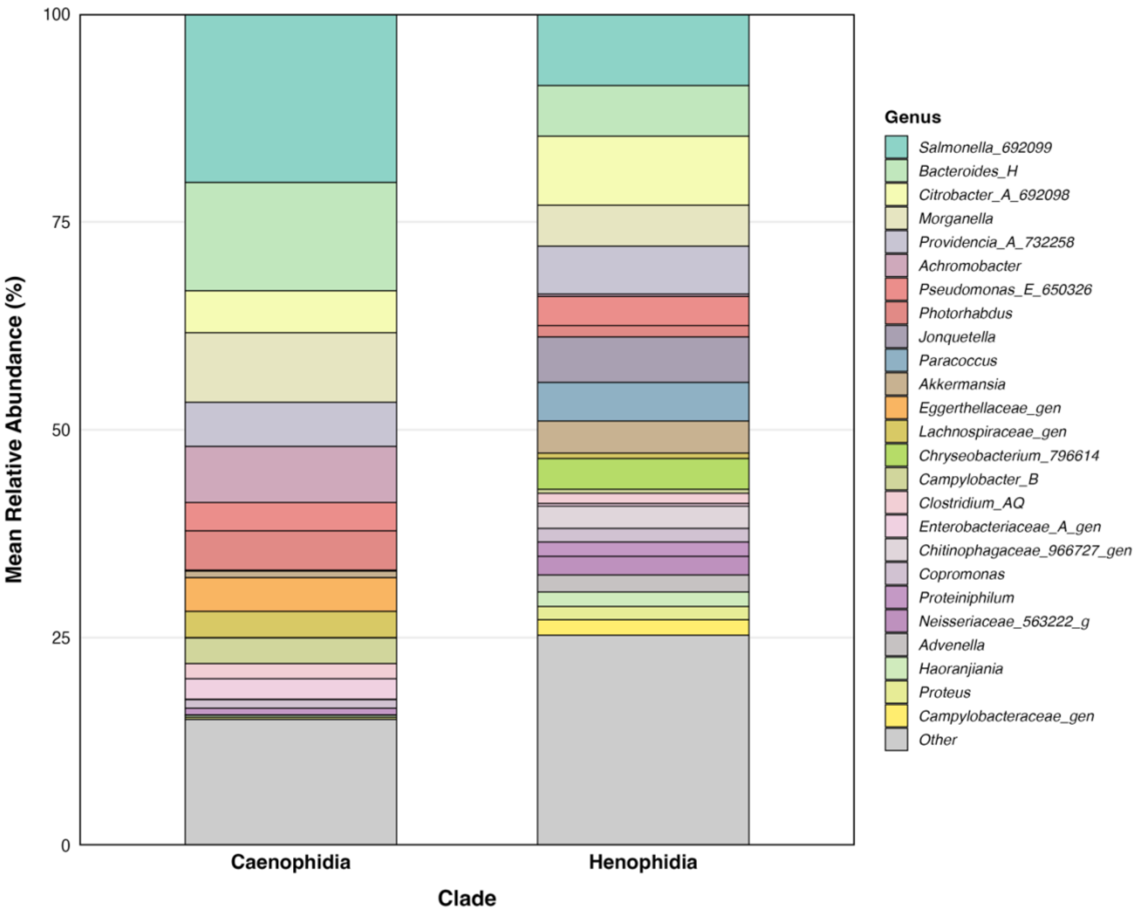


**Fig. S1.** Genus-level compositional profiles by clade. Stacked bar plots showing mean relative abundance of the 25 most abundant genera in *Caenophidia* and *Henophidia* cloacal microbiota. Each coloured segment represents a genus, with height proportional to mean relative abundance across all samples within that clade. Genera below the top 25 are grouped as "Other" (grey).

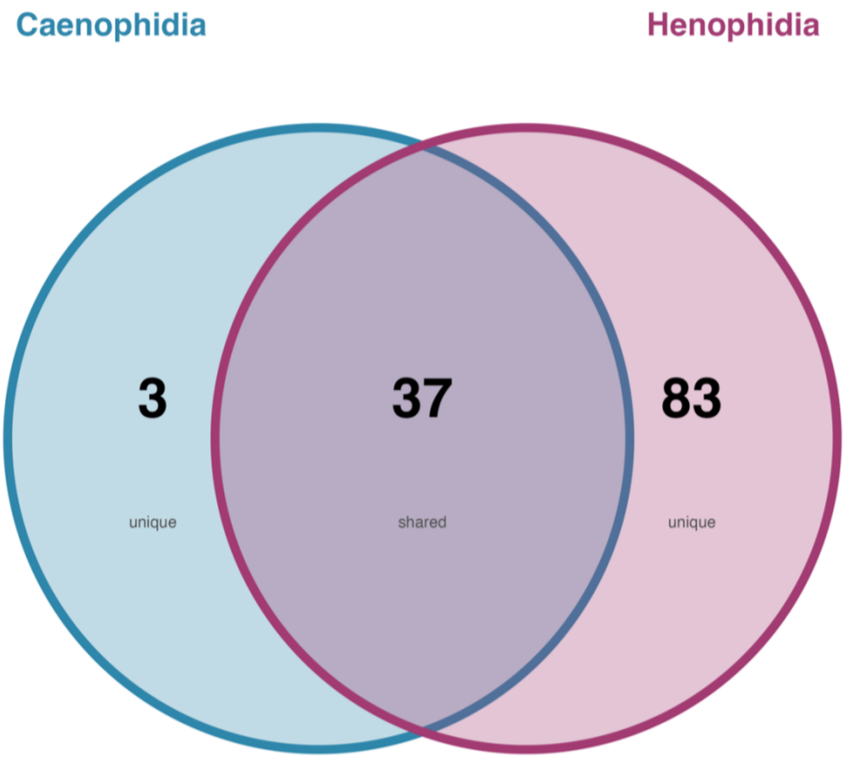


**Fig. S2.** Core and clade-specific genera distribution. Venn diagram showing the number of genera present in *Caenophidia*, *Henophidia*, or both clades (core). Genera were included if detected at ≥0.01% relative abundance in ≥20% of samples within at least one clade.

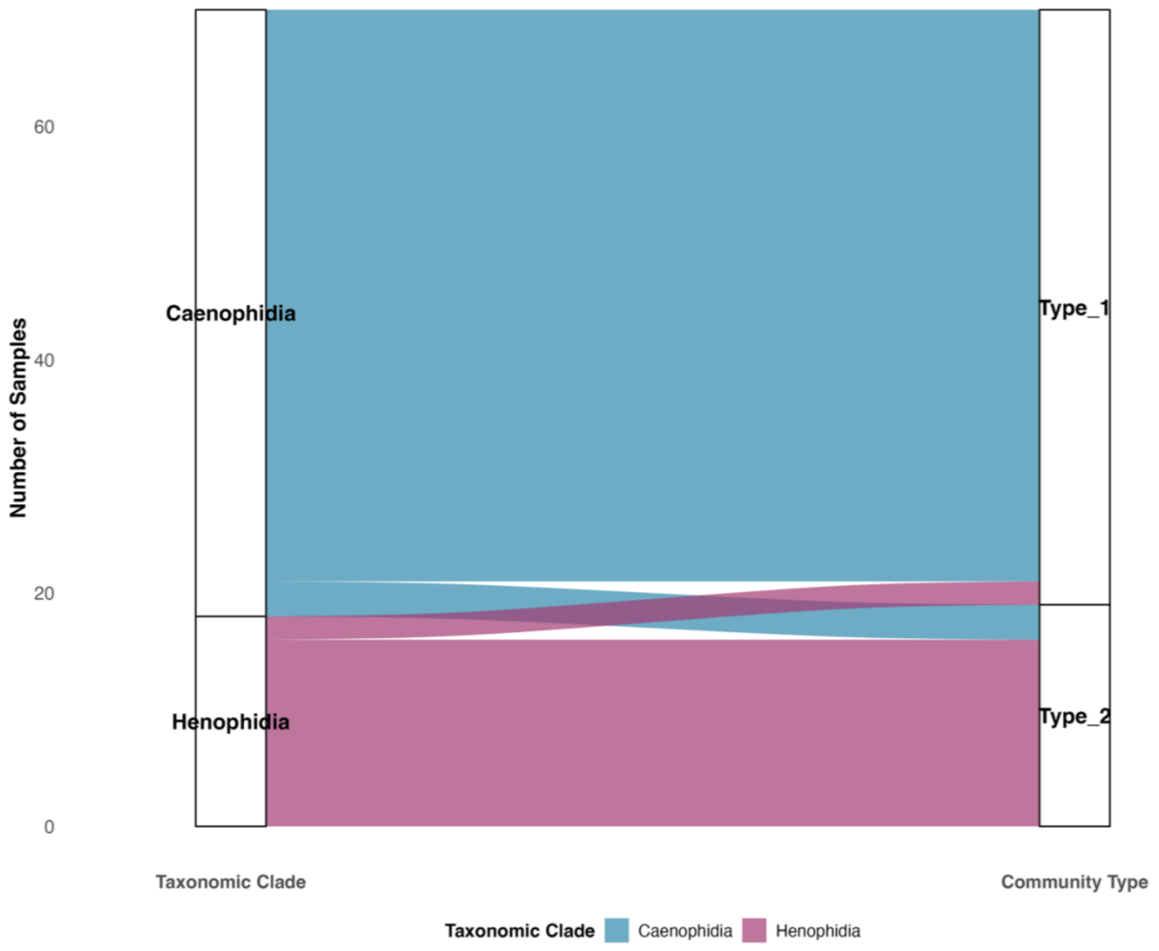


**Fig. S3.** Alignment of taxonomic clades and DMM community types. Alluvial diagram showing the distribution of samples from taxonomic clade (*Caenophidia*, *Henophidia*) to DMM defined community types (Type 1, Type 2). Ribbon width is proportional to the number of samples in each clade-community combination. Type 1 comprises 51 samples (49 *Caenophidia*, 2 *Henophidia*), whilst Type 2 comprises 19 samples (3 *Caenophidia*, 16 *Henophidia*).

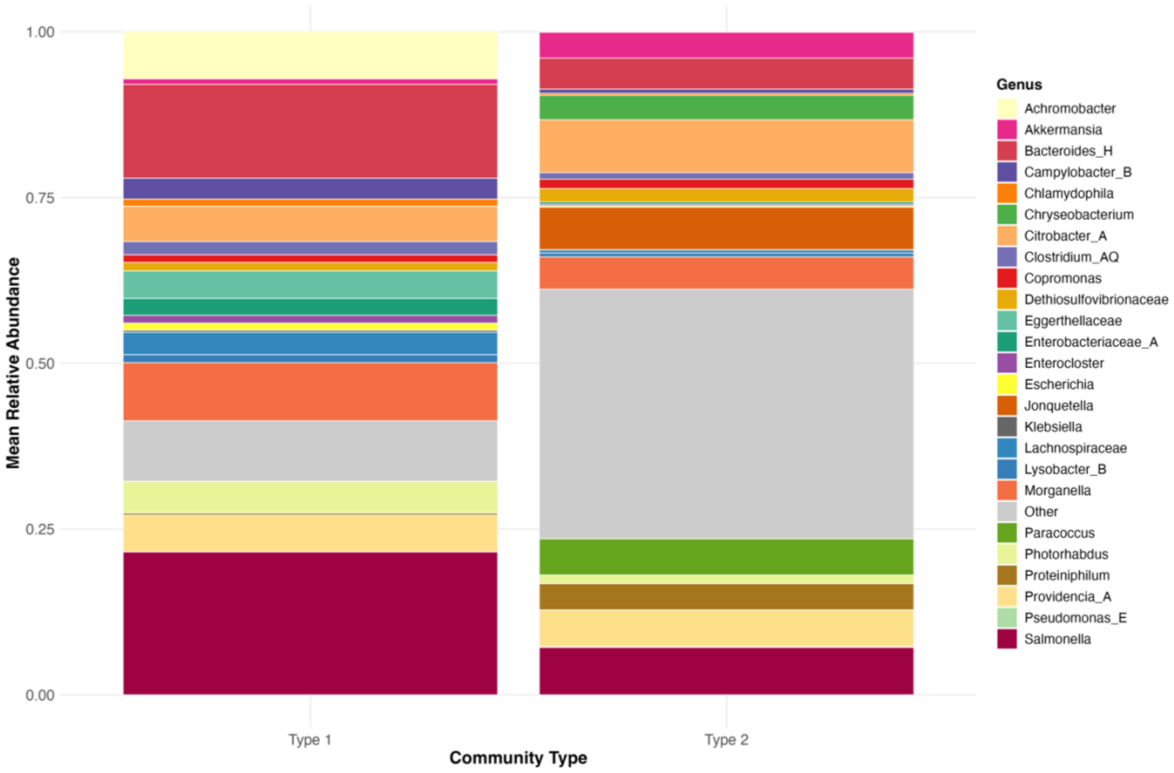


**Fig. S4**. Taxonomic composition of DMM community types. Stacked bar plot showing mean relative abundance of the top 25 most abundant genera across all samples for each community type, with remaining genera aggregated as "Other". Bar height represents 100% relative abundance. Type 1 (n = 51 samples) and Type 2 (n = 19 samples) display distinct compositional profiles, with differences in both the presence of specific genera and their relative abundances. Each coloured band represents a different bacterial genus, with band height proportional to mean relative abundance within that community type.

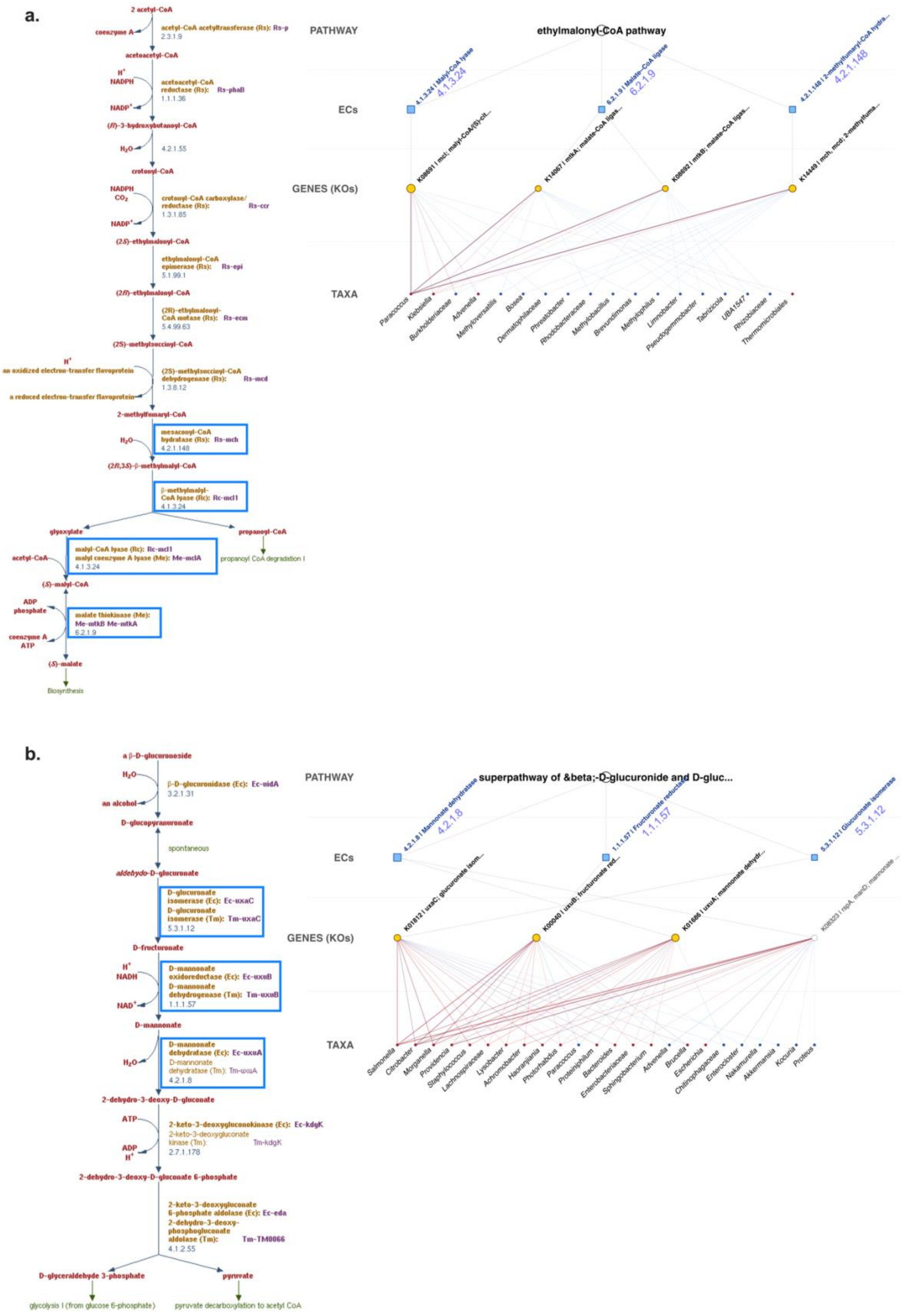
a.
2 acetyl-CoA
coenzyme A
acetyl-CoA acetyltransferase (Rs): Rs-p
2.3.1.9
acetoacetyl-CoA
acetoacetyl-CoA reductase (Rs): Rs-phaB
1.1.1.36
(R)-3-hydroxybutanoyl-CoA
4.2.1.55
crotonyl-CoA
crotonyl-CoA carboxylase/reductase (Rs): Rs-ccr
1.3.1.85
(2S)-ethylmalonyl-CoA
ethylmalonyl-CoA epimerase (Rs): Rs-epi
5.1.99.1
(2R)-ethylmalonyl-CoA
(2R)-ethylmalonyl-CoA mutase (Rs): Rs-ecm
5.4.99.63
(2S)-methylsuccinyl-CoA
an oxidized electron-transfer flavoprotein
(2S)-methylsuccinyl-CoA dehydrogenase (Rs): Rs-mcd
1.3.8.12
a reduced electron-transfer flavoprotein
2-methylfumaryl-CoA
mesaconyl-CoA hydratase (Rs): Rs-mch
4.2.1.148
(2R,3S)-β-methylmalyl-CoA
β-methylmalyl-CoA lyase (Rc): Rc-mcl1
4.1.3.24
glyoxylate
propanoyl-CoA
malyl-CoA lyase (Rc): Rc-mcl1
malyl coenzyme A lyase (Me): Me-mclA
4.1.3.24
propanoyl CoA degradation I
acetyl-CoA
(S)-malyl-CoA
ADP
phosphate
malate thiokinase (Me): Me-mtkB Me-mtkA
6.2.1.9
coenzyme A
ATP
(S)-malate
Biosynthesis
PATHWAY
ethylmalonyl-CoA pathway
ECs
4.1.3.24
6.2.1.9
4.2.1.148
GENES (KOs)
TAXA
Paracoccus
Klebsiella
Burkholderiaceae
Advenella
Methyloversatilis
Bosea
Dermatophilaceae
Phreatobacter
Rhodobacteraceae
Methylobacillus
Brevundimonas
Methylophilus
Limnobacter
Pseudogemmobacter
Tabrizicola
UBA1547
Rhizobiaceae
Thermomicrobiales
b.
a β-D-glucuronoside
β-D-glucuronidase (Ec): Ec-uidA
3.2.1.31
an alcohol
D-glucopyranuronate
spontaneous
aldehydo-D-glucuronate
D-glucuronate isomerase (Ec): Ec-uxaC
D-glucuronate isomerase (Tm): Tm-uxaC
5.3.1.12
D-fructuronate
NADH
D-mannonate oxidoreductase (Ec): Ec-uxuB
D-mannonate dehydrogenase (Tm): Tm-uxuB
1.1.1.57
D-mannonate
D-mannonate dehydratase (Ec): Ec-uxuA
D-mannonate dehydratase (Tm): Tm-uxuA
4.2.1.8
2-dehydro-3-deoxy-D-gluconate
ATP
2-keto-3-deoxygluconokinase (Ec): Ec-kdgK
2-keto-3-deoxygluconate kinase (Tm): Tm-kdgK
2.7.1.178
ADP
2-dehydro-3-deoxy-D-gluconate 6-phosphate
2-keto-3-deoxygluconate 6-phosphate aldolase (Ec): Ec-eda
2-dehydro-3-deoxy-phosphogluconate aldolase (Tm): Tm-TM0066
4.1.2.55
D-glyceraldehyde 3-phosphate
pyruvate
glycolysis I (from glucose 6-phosphate)
pyruvate decarboxylation to acetyl CoA
PATHWAY
superpathway of β-D-glucuronide and D-gluc...
ECs
4.2.1.8
1.1.1.57
5.3.1.12
GENES (KOs)
TAXA
Salmonella
Citrobacter
Morganella
Providencia
Staphylococcus
Lachnospiraceae
Lysobacter
Achromobacter
Haoranjiania
Photorhabdus
Paracoccus
Proteiniphilum
Bacteroides
Enterobacteriaceae
Sphingobacterium
Advenella
Brucella
Escherichia
Chitinophagaceae
Enterobacter
Nakamurella
Akkermansia
Kocuria
Proteus

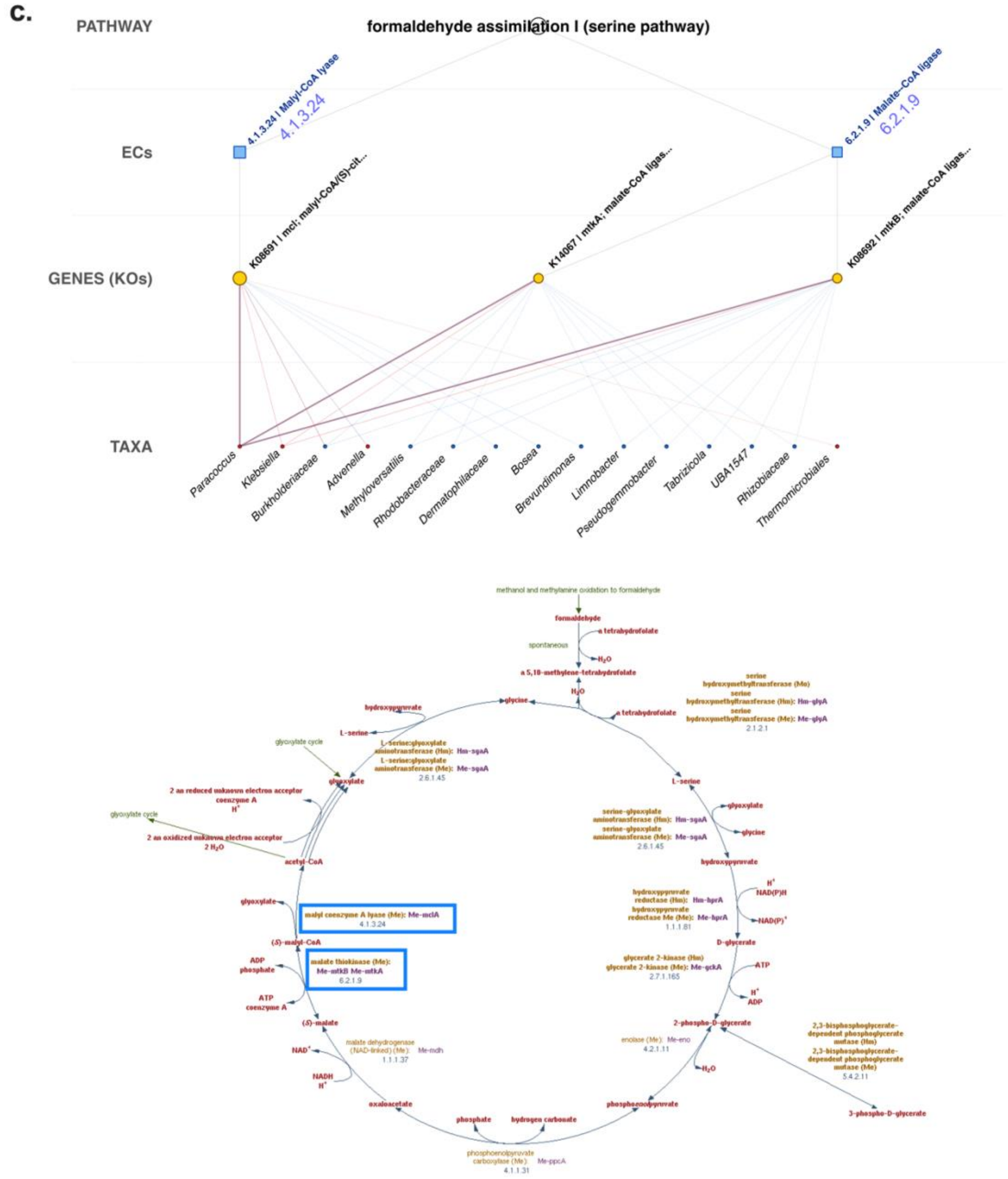


Fig. S5. Pathway–taxon linkage networks paired with reference metabolic pathway diagrams for three representative MetaCyc pathways. Three pathways are presented: (a.) ethylmalonyl-CoA pathway, (b.) superpathway of β-D-glucuronide and D-glucuronate degradation, and (c.) formaldehyde assimilation I (serine pathway). (a. and b.) The left panel and (c.) lower panel show the corresponding reference metabolic pathway diagram obtained from the MetaCycderived database hosted at the INRAe TrypanoCyc resource (http://vm-trypanocyc.toulouse.inra.fr/), in which enzymes identified in our linkage analysis are highlighted by blue boxes; Enzyme Commission (EC) numbers are shown in each highlighted reaction step to allow direct cross-referencing with the linkage network in the right panel. The right panel shows the four-tier linkage network, displaying the hierarchical mapping from pathway (top) through EC numbers (squares) and KEGG Orthologs (KOs, circles) to contributing genera (bottom). KO nodes highlighted in gold denote KEGG orthologs that were significantly differentially abundant between clades (adjusted $p < 0.05$, effect size $\geq 1.0$, CLR difference $> 1.0$); EC nodes in blue indicate enzyme activities. Transparent nodes correspond to non-significant ECs or KOs retained within the pathway definition. Edges connecting KOs to genera are coloured by clade enrichment (red: *Caenophidia*; blue: *Henophidia*), with edge width proportional to the relative contribution of each taxon. Node size is scaled by abundance. Cross-referencing the EC numbers between the two panels indicates the position of each differentially abundant enzyme within the canonical biochemical pathway.
